# Emergent simplicities in an individual cell’s stochastic response to disruptive change

**DOI:** 10.64898/2025.11.23.690018

**Authors:** Kunaal Joshi, Karl F. Ziegler, Shaswata Roy, Rhea Gandhi, Jack Stonecipher, Rudro R. Biswas, Srividya Iyer-Biswas

**Affiliations:** Department of Physics and Astronomy, Purdue University, West Lafayette, Indiana, USA; Weldon School of Biomedical Engineering, Purdue University, West Lafayette, Indiana, USA; Claremont McKenna College, Claremont, California, USA; Gustavus Adolphus College, St Peter, Minnesotta, USA; Santa Fe Institute, Santa Fe, New Mexico, USA

**Author notes:** These authors contributed equally to this work (Kunaal Joshi, Karl F. Ziegler).

## Abstract

Here we seek and find recurring patterns of behaviors in the stochastic response of an individual bacterial cell to a disruptive change in growth conditions. Building on the known scaling law that a single timescale, a cellular unit of time, governs stochastic growth and division of individual bacterial cells under constant growth conditions, here we show using experimental data that a dynamic rescaling of the cellular unit of time captures the predominant effect of temporal variations in environmental conditions. Furthermore, we identify the instantaneous exponential growth rate as the scaling factor that scales the internal clocks of the cells to the laboratory time. Our results reveal the natural representation for these time-dependent dynamics. When recast in its terms the cell age distribution for suitable initial conditions evolves under *time-invariant* rules even as growth conditions remain dynamic! Through the experimental realization at different temperatures of otherwise identical disruptive changes, we not only substantiate the general applicability of the cellular frame of reference but also uncover more emergent simplicities. Motivated by this representation, when time and instantaneous growth rate are expressed in terms of their naturally dimensionless counterparts, remarkably consistent patterns are revealed. These include a unimodal-bimodal-unimodal transition in the shape of the instantaneous growth rate distribution as cells initially in homeostasis experience a disruptive change in nutrient quality and subsequently recover and attain a new homeostasis. The remarkable scaling of the pattern of responses across temperatures suggests that the organizational rules and processes governing the response to disruptive change in nutrient quality remain the same at different temperatures. While the progression of the response appears to proceed at different tempos at different temperatures, upon shifting from the laboratory to the cellular frame of reference these changes progress at the same pace even at different temperatures.

---

Living organisms routinely experience temporal variations in environment and adjust their internal processes and organization accordingly. Are there recurring patterns in an individual bacterial cell’s physiological adaptation to temporal variations in external conditions which transcend system and environment specific details?

Approaching answering the question by directly performing high-precision quantitative livecell dynamic experiments on statistically identical cells subjected to a range of precisely controlled temporal variations, representative of what a typical organism may experience in nature, is currently impractical: these environments are inherently high-dimensional and an organism can have a large dynamic range of responses. In the absence of a clear separation of timescales, such as between the timescale of extrinsic variation in ambient conditions and the typical lifetime of the organism, the issues are further exacerbated (*1*). Even if such comprehensive datasets were available, systematic extraction of recurring patterns of behaviors, such as the emergent simplicities previously reported for cells in time-invariant conditions (*2–5*), is currently not feasible. Even for experiments involving the simplest unicellular organisms, such as bacteria, these challenges remain (*1, 6–11*).

## Results

### Emergent simplicity and a natural representation of cellular dynamics in time-varying conditions

Here we report the discovery of an emergent simplicity which captures the predominant effect of *temporal variations* in external conditions on the dynamics of individual bacterial cell growth and division. To contextualize the result, we first summarize the known emergent simplicity corresponding to *constant* growth conditions: A single timescale, a cellular unit of time, governs stochastic growth and division dynamics under constant growth conditions (*2–5*). In particular, the mean-rescaled interdivision time distributions from different temperatures, and the corresponding mean-rescaled cell age distributions, undergo scaling collapses (*2–5*).

Now consider the following proposal for generalization to dynamic conditions: *The net effect of the temporal variation in growth conditions is to simply rescale the cellular unit of time at each instant*. In other words, the time-dependence of this internal cellular timescale encapsulates the effects of external variations in ambient conditions on the growth and division of individual cells. As we show below, this proposal is compellingly borne out experimentally.

Our proposal also motivates *a natural representation* for cellular dynamics in time-varying conditions: the key insight is to use the natural dimension-free rescaled variables, transforming from the absolute ‘lab time’, *t*, and absolute lab-measured age, *τ*, of the cell to the corresponding quantities, (*t*_*r*_, *τ*_*r*_), measured using the ‘internal’ clocks of the cells:

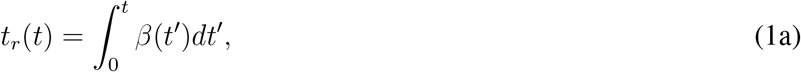

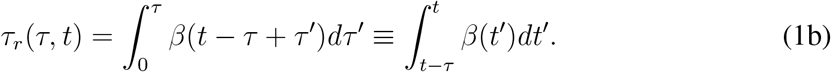

Requiring that the temporal scaling factor *β*(*t*) always remain positive, these equations uniquely specify (*t*_*r*_, *τ*_*r*_) given (*t, τ*).

Furthermore, the implication of the existence of a single (dynamic) governing timescale is that the division propensity, in terms of the intrinsic clock of the cell, is independent of explicit dependence on absolute lab time: *α*_*r*_(*τ*_*r*_) = *α*(*t, τ*)*/β*(*t*). It follows that the division time distributions from different constant growth conditions, once mean-rescaled, should undergo a scaling collapse, as is indeed observed in experimental data (*2*). The rescaled propensity, *α*_*r*_(*τ*_*r*_), can be expressed in terms of the (static) division time distribution in the intrinsic cellular frame of reference: 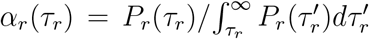. Upon changing variables, we find that the age distribution in the cellular frame is related to the age distribution in the lab frame as:

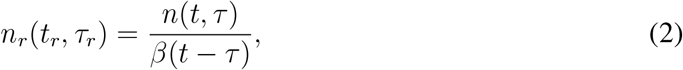

where *t, τ* are related to *t*_*r*_, *τ*_*r*_ through Eq. (1). Physically, Eq. (2) implies that to obtain the age distribution in the cellular frame, we need to scale the age distribution in lab frame by the temporal scaling factors at the corresponding times of birth.

### Experimentally relevant case of special interest

To motivate convenient experimental design, we consider the special case in which the system starts with steady-state age distribution, then experiences a change in conditions, leading eventually to a different steady state: In this case, to begin with the age distribution in the cellular reference frame is already in steady state (i.e., independent of *t*_*r*_), and continues to be time invariant even as the growth conditions change. All changes in this case are simply encapsulated in the scaling factor *β*, while the cellular reference frame always behaves as if it were in constant growth conditions. There is practical interest in the case when population size is held constant: the mother cell is retained at each division and the daughter cell(s) removed from the experimental arena, resulting in a non-growing constant population number of cells in the experiment. Single-cell technologies such as the SChemostat and the Mother Machine facilitate the experimental realization of this scenario (*2, 12–16*). These experiments also allow the direct tracking of the age distribution as a function of (lab) time. Therefore, if the population size is also held constant, the cell age distribution in the cellular reference frame is then given by,

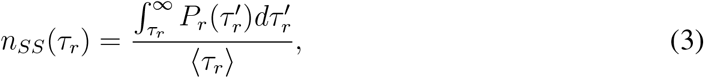

where, *P*_*r*_ is the division time distribution in the cellular reference frame. We introduce the average division time in the cellular reference frame, ⟨*τ*_*r*_⟩ = *τ*_*r*_*P*_*r*_(*τ*_*r*_)*dτ*_*r*_, to normalize *n*_*SS*_(*τ*_*r*_) to 1. The age distribution in the lab frame can now be obtained from Eq. (2):

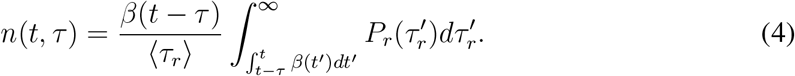

Given that the system was initially in steady state, we can obtain *P*_*r*_ from the initial steady state division time distribution *P*_*s*_ in the lab frame, and the initial steady state scaling factor *β*_0_, using the relation *τ*_*r*_ = *β*_0_*τ* in steady state. Thus,

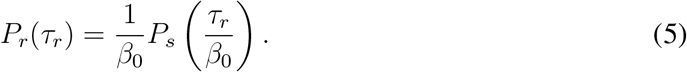

### Experimental realization via SChemostat technology of statistically-identical non-interacting individual cells being subjected to a precisely controlled disruptive change

The SChemostat technology (*2*) is uniquely suited to performing high-precision high-throughput experiments on statistically-identical non-interacting individual bacterial cells subjected to precisely controlled temporal variations in environmental conditions. Here we focus on the experimental scenario in which an asynchronous population of cells is initially in steady state in constant conditions (complex nutrient media) and then undergo an abrupt shift to a novel condition (minimal nutrient media), which they have never previously encountered in their (multigenerational) lifetimes. We observe these cells for tens of generations after the abrupt change in conditions—long enough for each cell to attain a new steady-state (Fig. 1). Since the initial population is asynchronous, different cells experience the switch in conditions at different ages (since the last division event). While at first glance the resulting trends may appear to display bewildering complexity, we are nevertheless able to extract from these data the following emergent simplicity.

**Figure 1.**
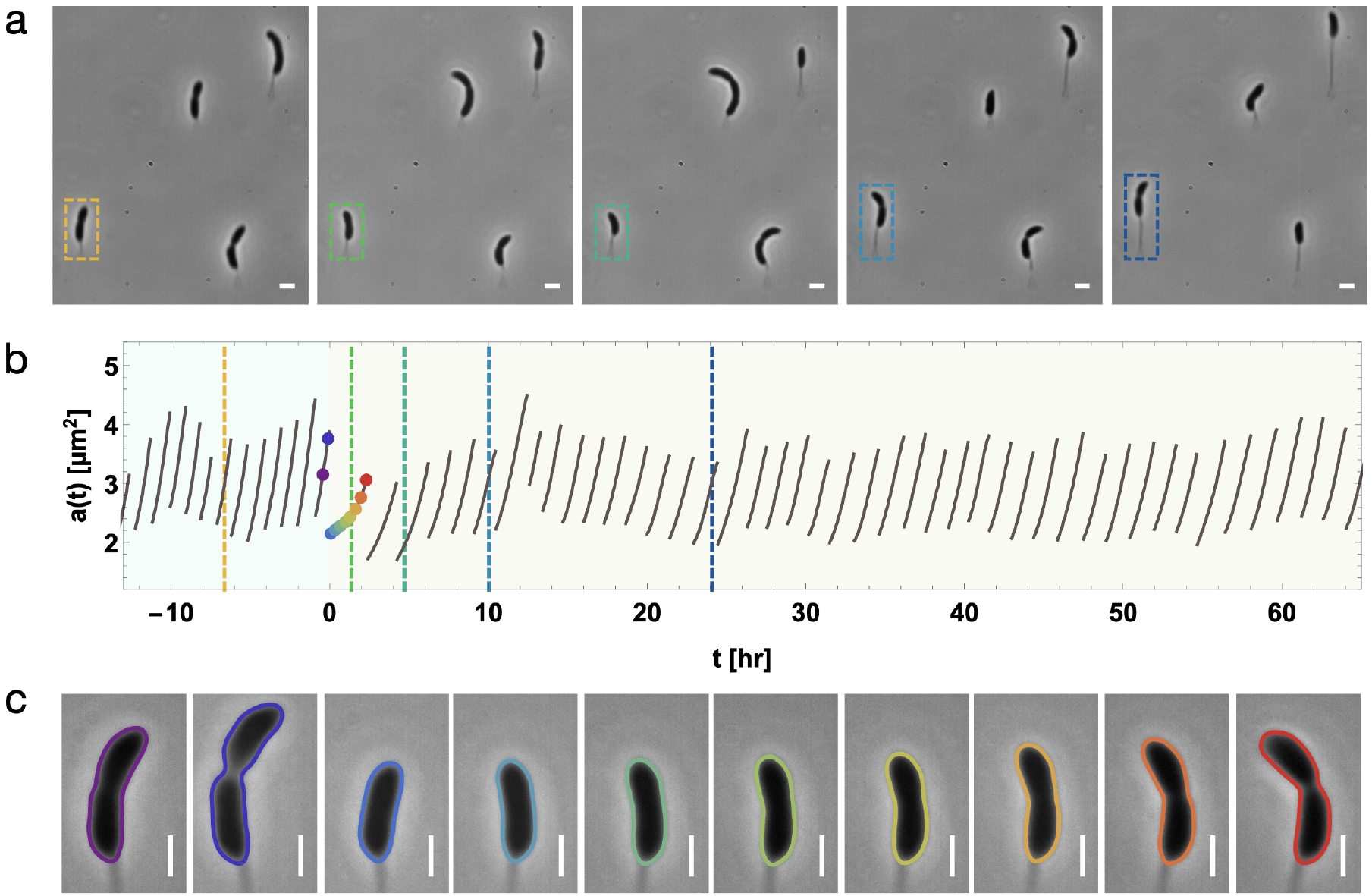
High-precision imaging of an individual cell’s stochastic response to disruptive change in nutrient quality performed using the SChemostat technology. **(a)** Snapshots of a typical field of view at five different time points in the SChemostat setup, containing four cells where the dashed box highlights the same cell with colors corresponding to the times indicated by dashed lines on the trajectory in (b). **(b)** The time evolution of the experimentally measured area growth trajectory of the highlighted cell in (a), as the growth media is abruptly switched from complex (light blue background) to minimal (light yellow background), with the switch occuring at *t* = 0. **(c)** Zoomed-in images of the highlighted cell in (b), superimposed with splines calculated by automated analysis, at approximately equally spaced time points chosen from generations immediately preceding and succeeding the switch in growth conditions. These times are indicated by points with corresponding colors in (b). Scalebar corresponds to 1*µ*m. Using these images and the equal time-spacing between images within each generation, the significant slowing down of cell growth immediately after the switch is visually evident.

### Emergent simplicity: The instantaneous exponential growth rate robustly encapsulates the dynamic rescaling of the cellular unit of time

First we extract the instantaneous growth rate: the time derivative of the logarithm of the cell size (*17–20*). Under time-invariant growth conditions the scaling factor that scales the internal timescale to match the external (or lab) time, *β*, has evidently been identified as proportional to the exponential growth rate of individual bacterial cells (*2*). Here, we provide experimental evidence to demonstrate that under *time-varying* growth conditions, the instantaneous exponential growth rate at a given instant in time (*t*) is proportional to the scaling factor at that instant in time (*β*(*t*)) (see Fig. 2 – when the age distributions are rescaled using the instantaneous growth rate as the scaling factor *β*(*t*), the distributions become invariant in time under all growth conditions tested). Thus the challenge in extracting dimension reduction from these complex and stochastic dynamics is conveniently overcome.

**Figure 2.**
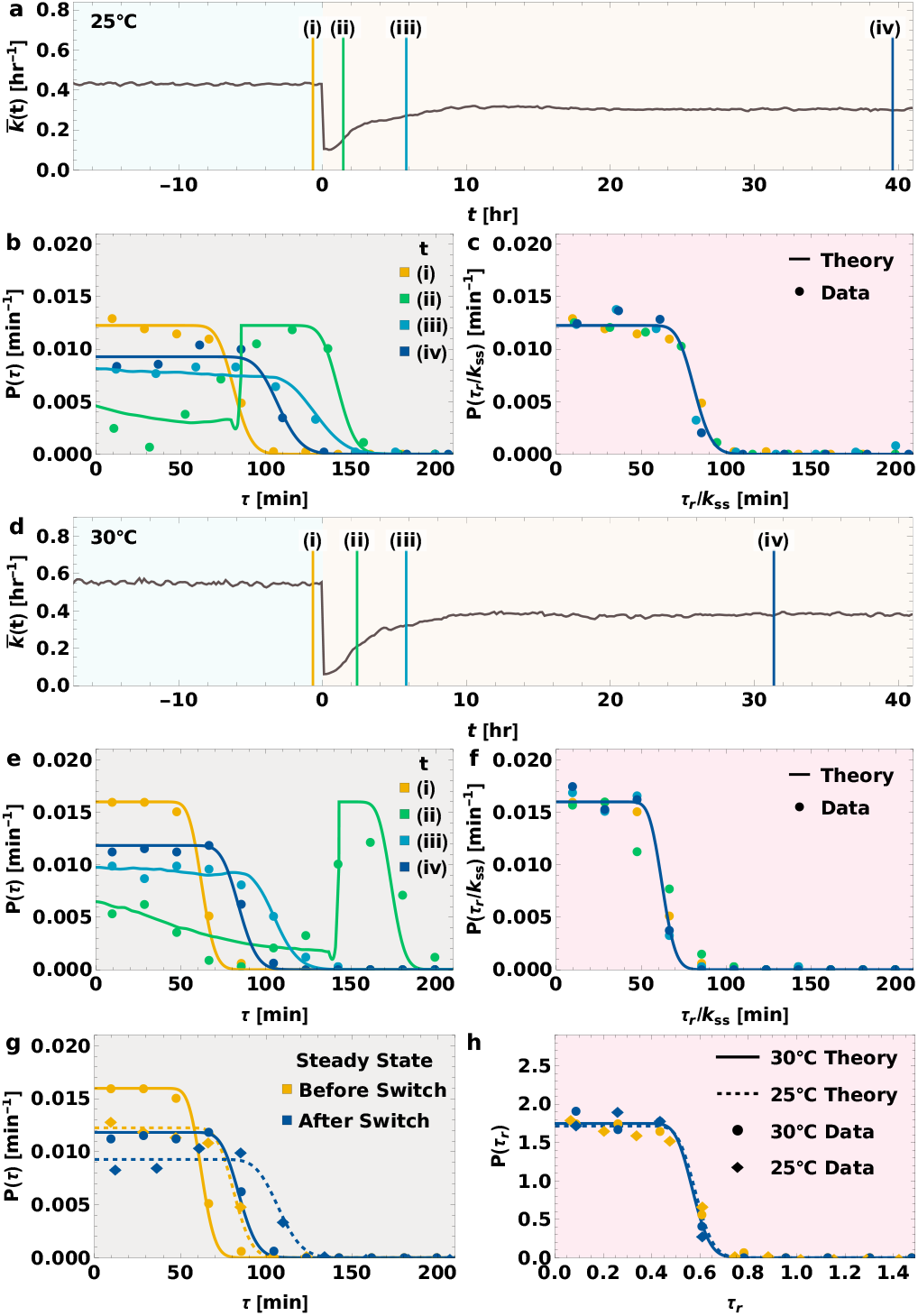
Emergent Simplicity: Dynamical rescaling by the instantaneous growth rate renders the dynamics time-invariant in the cellular frame of reference. **(a)** The mean of the asynchronous instantaneous growth rate is plotted across the abrupt switch in growth conditions, from complex to minimal media at 25^°^ C. This mean value is used along with the steady state division time distribution to predict the transient age distributions without any free parameters. **(b)** Distributions of cell ages at different experiment time instants corresponding to (i–iv) marked in (a) with colored lines. The points represent the experimentally measured distributions, while the solid lines are the theoretical predictions given by Eq. (4). **(c)** The age distributions in the cell’s intrinsic frame of reference (*τ*_*r*_) are plotted at the corresponding time instants. These distributions remain invariant in time, and match the theoretical prediction given by Eq. (3). These distributions are scaled by a constant value (*k*_*ss*_, or the average steady-state growth rate before switch) to aid in visual comparison. **(d–f)** The corresponding plots for the same experiment conducted at 30°C. **(g–h)** A comparison of the steady state age distributions before and after the instantaneous switch in growth conditions at the two different temperatures. The distributions in the lab frame of reference (g) are distinct, but they all overlap in the cell’s intrinsic frame of reference (h).

### Emergent simplicity: In the cellular frame of reference the cell age distribution remains time invariant even after disruptive change, for cells initially in a balanced growth condition

Upon rescaling the time in the lab frame by the population mean instantaneous exponential growth rate (previously identified as proportional to the the scaling factor *β*(*t*)) through Eq. (1), we find that the cell-age distributions sampled at different times since the switch, which differ dramatically in shape as they adjust from the previous steady-state to the new conditions, all collapse onto the same time-invariant distribution in the cellular reference frame (Fig. 2). This observation validates that (i) the scaling factor is indeed proportional to the instantaneous exponential growth rate (there is freedom in choosing the constant of proportionality), and (ii) in the cellular reference frame, the cell indeed behaves as if it were under constant growth conditions, with all temporal changes in growth conditions encapsulated in the scaling factor. Furthermore, the predicted age distributions for both the cellular reference frame (Eq. (3)) and lab frame (Eq. (4)) match the experimental data without the need for any fitting parameters (Fig. 2).

### Emergent simplicity: Scaling across temperatures of cellular responses to disruptive change in nutrient quality

Taking further advantage of the unique capabilities of the SChemostat technology, to test the generality of the results reported above, we performed similar high-precision high-throughput experiments on statistically-identical non-interacting individual bacterial cells subjected to precisely controlled temporal variations in nutrient conditions, now at a different temperature: previously at 30°C and now at 25°C. For instantaneous switch from complex to minimal media at 30°C, we see an instantaneous sharp decline in growth rate almost all the way to zero, followed by a gradual exponential recovery to the new steady state value which is smaller than the initial steady state value. For the same disruptive change, though now at 25°C, we see a similar response, except the steady state values are scaled by a constant factor (Fig. 3 b), which is consistent with the Arrhenius scaling law (*2, 3*). *Our results thus reveal the natural representation for these time-dependent dynamics*.

**Figure 3.**
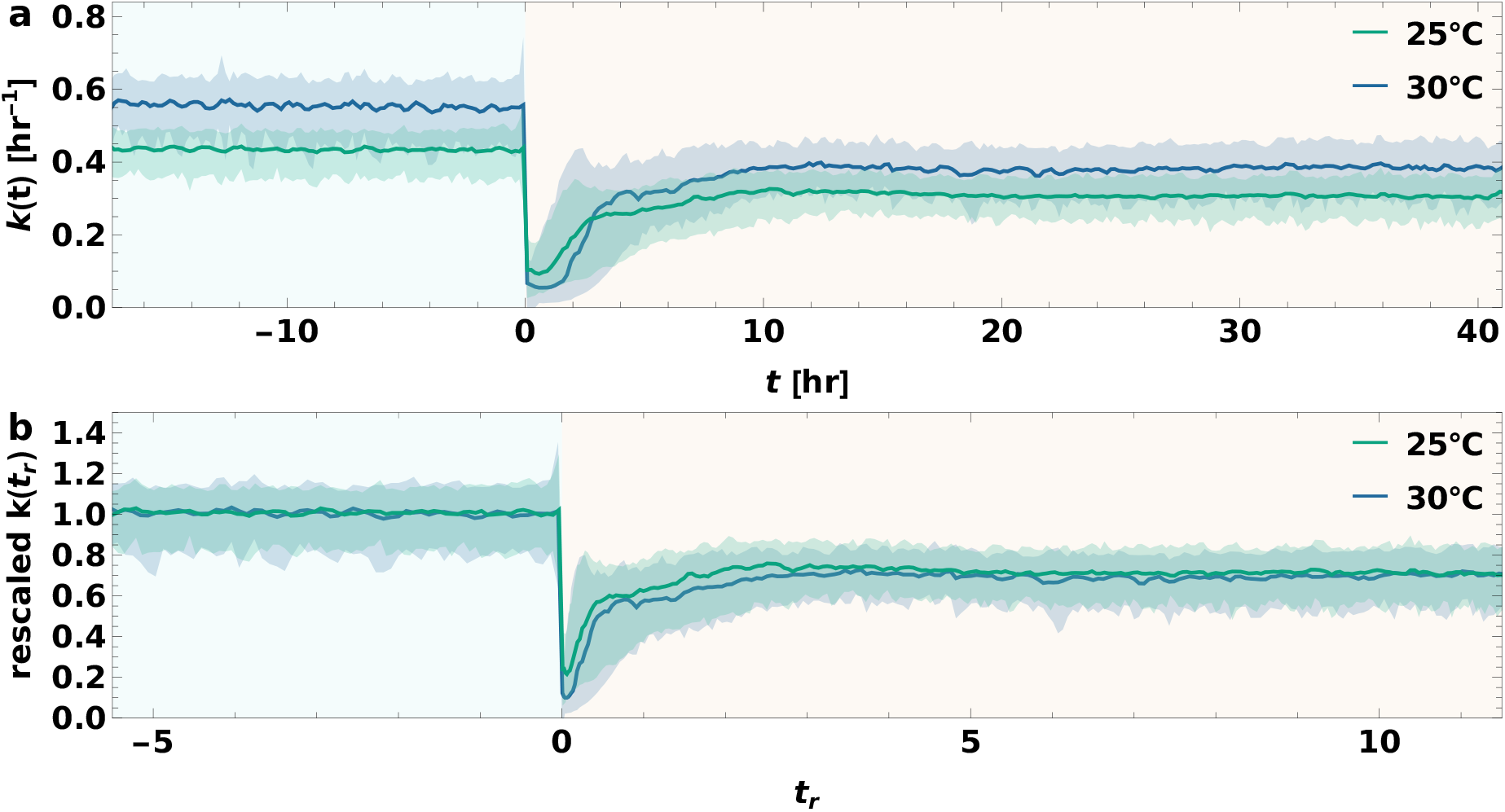
The instantaneous growth rate of cells subjected to an abrupt shift in nutrient quality at different temperatures in the laboratory and cellular frames of reference. **(a)** The median (solid line) and the 90% quantile range (shaded region) of the asynchronous instantaneous growth rate is plotted as a function of time across the abrupt switch (at *t* = 0) in growth conditions from complex to minimal media at 25°C (green) and 30°C (blue). **(b)** The instantaneous growth rates in (a) are rescaled by their mean steady state values prior to switch, and plotted as a function of the time in cell’s intrinsic frame of reference given by Eq. (1). The rescaled steady state instantaneous growth rate values after switch overlap.

### Emergent simplicities beyond the temporal scaling ansatz: consistent pattern across different temperatures of transient bimodality in the instantaneous growth rate distributions immediately following disruptive change in nutrient quality

Notably, upon being subjected to disruptive change, the distribution of instantaneous growth rates transitions from the initial to the final steady state forms, which are tight unimodal distributions, through intermediate *transient bimodal distributions*. Motivated by the natural representation, when time and instantaneous growth rate are expressed in terms of their naturally dimensionless counterparts, remarkable similarity in the progression of this unimodal-bimodal-unimodal transition is observed at both 25°C and 30°C (Fig. 4). Immediately following the change in conditions the growth-rate distribution becomes bimodal with a strong mode at lower values immediately after the switch. Thereafter, locations of the modes remain approximately constant, but the weight gradually shifts from the lower to the higher mode, suggesting a binary rather than a graded response. *The remarkable scaling of the pattern of responses across temperatures suggests that the organizational rules and processes governing the response to disruptive change in nutrient quality remain the same at different temperatures. While the progression of the response appears to proceed at different tempos at different temperatures, upon shifting from the laboratory to the cellular frame of reference these changes progress at the same pace even at different temperatures*.

**Figure 4.**
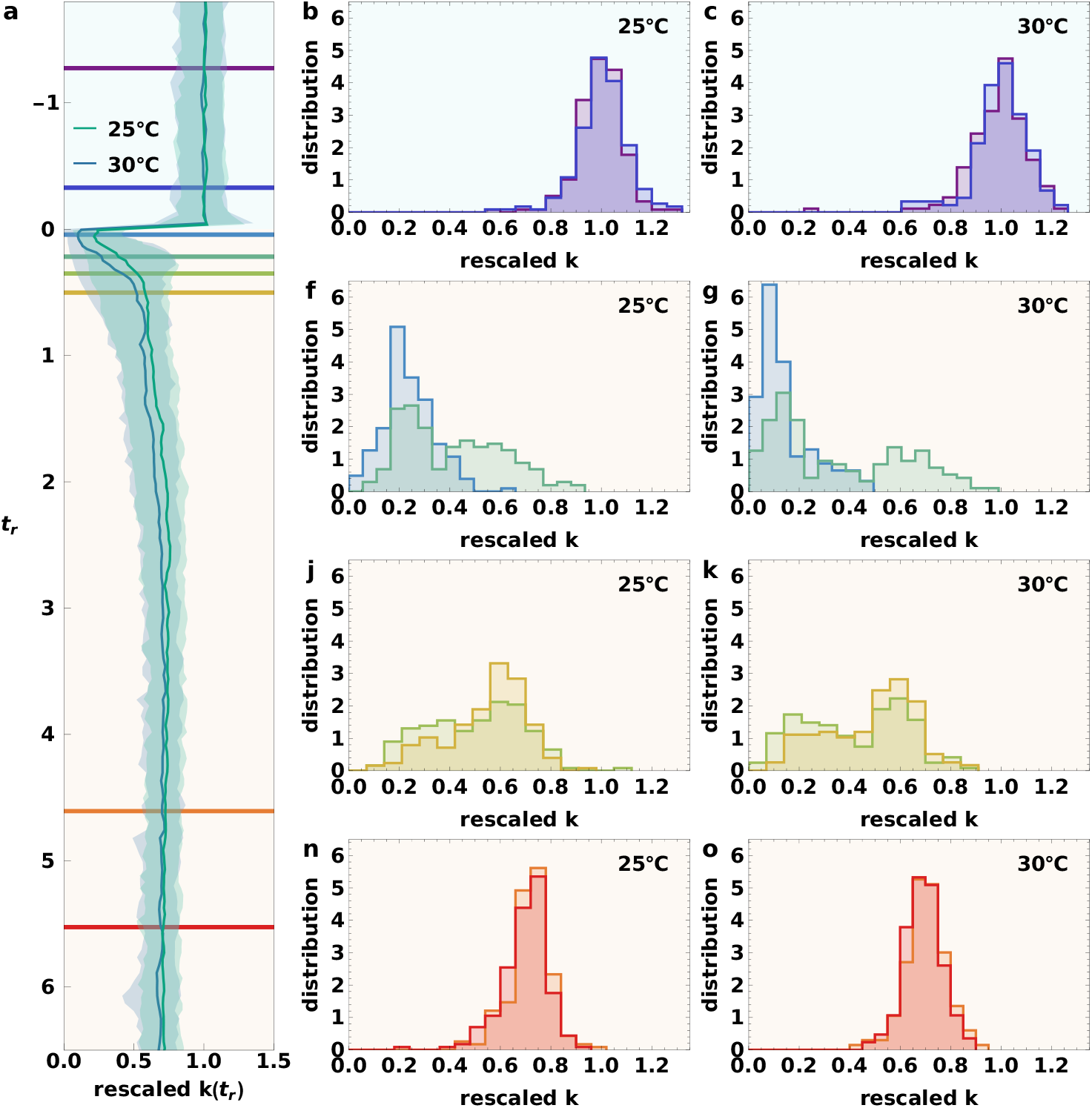
The pattern of unimodal-bimodal-unimodal transition in the shape of the rescaled instantaneous growth rate distribution, as cells initially in homeostasis experience a disruptive change in nutrient quality and subsequently recover and attain a new homeostasis, is remarkably consistent across temperatures. **(a)** The median along with the 90% quantile interval of the instantaneous growth rate rescaled by the pre-switch steady state mean is plotted as a function of time in the cell’s itrinsic frame of reference, across the abrupt switch in growth conditions from complex to minimal media at 25°C (green) and 30°C (blue). **(b, f, j, n)** The instantaneous growth-rate distributions at 25°C, rescaled by the pre-switch steady state mean, are plotted at different rescaled times after switch, corresponding to the time points in the cellular frame of reference marked in (a) by lines of corresponding colors. The final steady-state distributions have a larger coefficient of variation than the initial ones. We see bimodality in the transient distributions, and as time progresses, the height of the mode at smaller *k* decreases along with a corresponding increase in the height of the other mode at larger *k*. **(c, g, k, o)** The corresponding rescaled *k* distributions at 30°C also show a remarkably consistent pattern with the distributions at 25°C, including the unimodal-bimodal-unimodal transition.

## Concluding remarks

We have presented a framework to characterize the time-varying aspects of cell growth and division dynamics under time-varying growth conditions through a dynamic rescaling of the cellular unit of time, and shown that the scaling factor is the time-varying instantaneous growth rate. This framework serves as a useful starting point for extending to time varying conditions aspects of cell growth and division dynamics which have been previously been well studied in constant growth conditions, such as the problem of cell size homeostasis (*12, 21–23*).

The mechanism underlying the observed behavior of transient bimodality in the adaptation of instantaneous growth rate to an instantaneous switch in growth conditions has yet to be explicated. The phenomenon of a shift in weight between modes cannot be explained simply through some mechanism leading to a high correlation in instantaneous growth rates across generations, since that would result in both modes shifting to the right with the same weights rather than the weight shifting between modes. Our preliminary investigations reveal that cell division may play a pivotal role in this shifting of the weight. This finding may serve as a useful clue for this promising line of future inquiry.

## Author Contributions

R.R.B. and S.I.-B. conceived of, designed and implemented the theoretical framework; K.J., S.R., R.R.B. and S.I.-B. performed analytic calculations; K.J. performed numerical computation under the guidance of S.I.-B.; K.F.Z. designed and refined experimental protocol under the guidance of S.I.-B.; K.F.Z. performed experiments with contributions from S.I.-B.; K.J. performed data analyses under the guidance of S.I.-B.; K.J., K.F.Z., S.R., R.G. and J.S. contributed to manual supervision of image analysis; K.J., R.R.B., and S.I.-B. wrote the paper with input from K.F.Z. and S.R.; S.I.-B. helped shape the research and analysis and supervised all aspects of the research, and saw it through to fruition.

## Acknowledgements

We thank Purdue University Startup funds, Purdue Research Foundation, the Purdue College of Science Dean’s Special Fund, and the Showalter Trust for financial support. K.J. and S.I.-B. acknowledge support from the Ross-Lynn Fellowship award. S.I.-B. thanks the Purdue Physics and Astronomy REU program for hosting and providing financial support for R.G. and J.S’s contributions. We thank the Iyer-Biswas group members for useful discussions. We are grateful to Charles Wright for insightful discussions and detailed feedback on the manuscript.

## Data and materials availability

Data reported here will be made available through a public repository along with this publication.

